# Endogenous memory reactivation during sleep in humans is clocked by slow oscillation spindle complexes

**DOI:** 10.1101/2020.09.16.299545

**Authors:** Thomas Schreiner, Marit Petzka, Tobias Staudigl, Bernhard P. Staresina

## Abstract

Sleep is thought to support memory consolidation via reactivation of prior experiences, with particular electrophysiological sleep signatures (slow oscillations (SOs) and sleep spindles) gating the information flow between relevant brain areas. However, empirical evidence for a role of endogenous memory reactivation (i.e., without experimentally delivered memory cues) for consolidation in humans is lacking. Here, we devised a paradigm in which participants acquired associative memories before taking a nap. Multivariate decoding was then used to capture endogenous memory reactivation during non-rapid eye movement (NREM) sleep in surface EEG recordings. Our results reveal reactivation of learning material during SO-spindle complexes, with the precision of SO-spindle coupling predicting reactivation strength. Critically, reactivation strength (i.e. classifier evidence in favor of the previously studied stimulus category) in turn predicts the level of consolidation across participants. These results elucidate the memory function of sleep in humans and emphasize the importance of SOs and spindles in clocking endogenous consolidation processes.

## INTRODUCTION

How do we strengthen memories while we sleep? The prime vehicle of systems consolidation is thought to be the reactivation of information encoded during prior wakefulness ^1–4^. Through reactivation, memory representations are relayed between the hippocampus and neocortical long-term stores, transforming initially labile representations into long-lasting memories during sleep ^5,6^. The communication between the hippocampus and neocortical networks is thought to be facilitated by an intricate interplay of the cardinal NREM sleep-related oscillations, namely cortical slow oscillations (SOs), thalamo-cortical sleep spindles and hippocampal sharp-wave ripples 7–12. SOs reflect fluctuations of the membrane potential and orchestrate transitions from neuronal silence (‘hyperpolarization’, i.e., down-state) to neuronal excitation (depolarization, i.e., up-state)^13,14^. Importantly, they initiate time windows of excitability and inhibition not only in cortical but also in subcortical areas ^15–17^. They trigger the emergence of sleep spindles in the thalamus ^18^, which nest in the excitable up-states of the SOs. Spindles have been shown to gate Ca^2+^ influx into dendrites, thereby facilitating synaptic plasticity ^19,20^. Importantly, recent evidence from two-photon imaging in mice suggests that Ca^2+^ influx is strongly amplified when spindles coincide with SO up-states ^21^. Lastly, hippocampal ripples are transient network oscillations and have been closely linked to reactivation/replay of learning experiences ^22,23^. They have been shown to occur in the excitable troughs of the spindle, suggesting that spindles might facilitate information transfer from the hippocampus to neocortical target sites ^24,25^. The efficacy of systems consolidation through memory reactivation might thus hinge on concurrent SO-spindle coupling, ensuring optimal conditions to ignite structural changes in cortical target sites ^8,11,26,27^.

Indeed, recent work in humans has revealed a key role of SO-spindle coupling during NREM sleep for behavioral expressions of consolidation. For instance, the precision of SO-spindle coupling, i.e., the exact timing of spindle maxima with respect to the SO up-state, has been shown to correlate with retention of declarative learning material ^28,29^. Moreover, levels of SO-spindle coupling track the rise and decline of memory performance across development ^30–32^. What is unknown, however, is whether there is a link between SO-spindle coupling and physiological expressions of consolidation, i.e., memory reactivation. A recent rodent study revealed that precise SO-spindle coupling is key for maintaining the reactivation of neural ensembles ^33^, but whether and how this relates to episodic memory consolidation in humans is unclear.

In humans, the study of memory reactivation during sleep has mainly relied on targeted memory reactivation (TMR) protocols ^34,35^. This experimental technique follows the rationale that reminder cues are presented during sleep to exogenously trigger memory reactivation. Intriguingly, presenting auditory reminder cues during NREM sleep reliably induces SO-spindle complexes ^36–38^. However, to what extent TMR-induced processes reflect natural/endogenous consolidation processes remains unknown.

Building on the work summarized above, we propose that SO-spindle complexes might clock endogenous memory reactivation in service of consolidation during human sleep. To test this notion, we devised an experimental paradigm in which participants acquired associative memories before taking a nap. Multivariate decoding was then used to assess endogenous memory reactivation during NREM sleep. In this work, we show that memory reactivation is specifically bound to the presence of SO-spindle complexes, with the precision of their coupling correlating with reactivation strength. Reactivation strength in turn predicts the extent of consolidation across participants. These findings elucidate the memory function of sleep in humans and illustrate the importance of SO-spindle coupling for clocking endogenous consolidation processes.

## RESULTS

Twenty participants (age: 20.75 ± 0.35; 17 female) took part in two experimental sessions. In both sessions they performed an episodic learning task, with memory performance being assessed before and after taking a 120 minutes nap (Fig. 1a). Depending on the experimental session, participants learned to associate verbs with images of objects or scenes during the pre-sleep learning phase. These stimulus categories were chosen as they recruit distinctive brain networks (e.g., lateral occipital complex for objects, parahippocampal place area for scenes ^39,40^), thus facilitating the analytical readout of endogenous, experience-dependent memory reactivation during sleep. Specifically, learning-related memory reactivation during sleep would manifest as enhanced representational evidence for the stimulus category learned before sleep (i.e., greater evidence for object representations after word-object encoding and greater evidence for scene representations after word-scene encoding, respectively).

**Figure 1.**
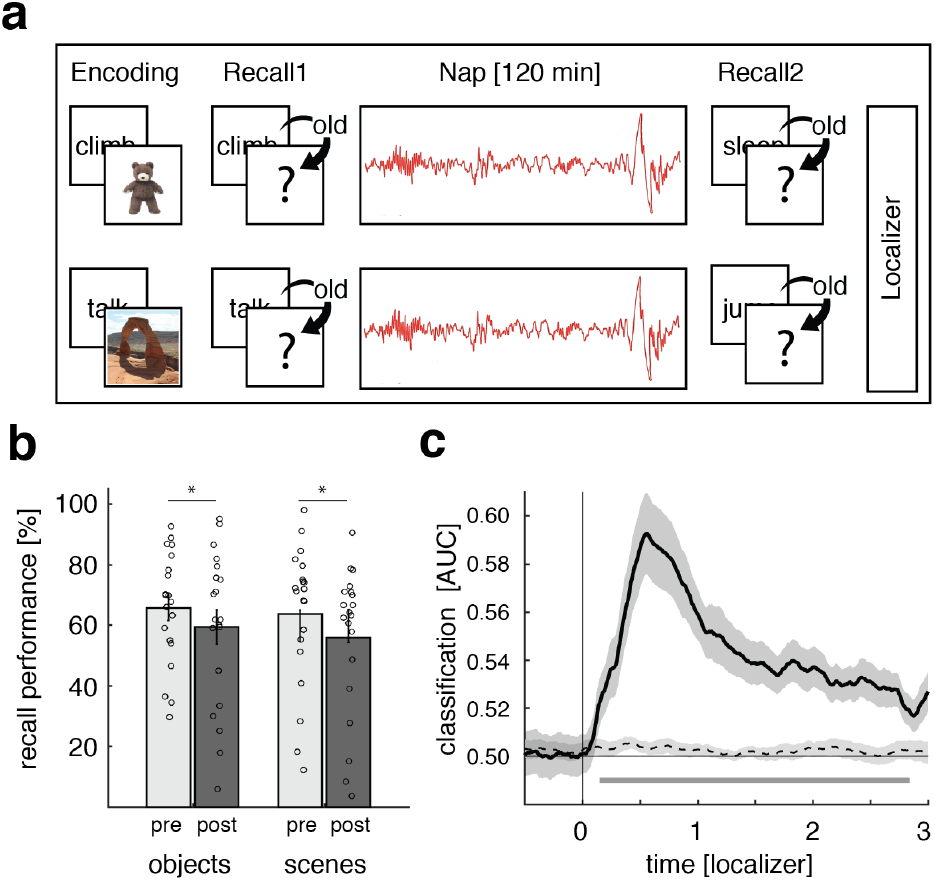
Experimental procedure, behavioral results & localizer task. (a) During encoding, participants were presented with 120 verb-object or verb-scene combinations (depending on experimental session). Memory performance was tested before and after a 120 minutes nap period. At the end of each session, participants performed a localizer task in which they processed a new set of object and scene images. (b) Behavioral results for both experimental sessions pre-(light gray) and post-sleep (dark gray). Bar graphs show mean (±SEM) percentage of recalled image exemplars out of correctly recognized verbs. Dots indicate individual memory performance of participants (N = 20). Stars denote significant differences as derived from a repeated measures ANOVA (p = 0.001). (c) Stimulus categories (objects vs. scenes) could be reliably decoded (above chance) from the localizer EEG data, starting around 150 ms post stimulus onset (the black solid line indicates decoding performance (±SEM)). The horizontal dashed line indicates surrogate decoding performance, which was estimated by shuffling the training labels 250 times. The vertical solid line indicates stimulus onset (time = 0). The lower horizontal gray line shows the temporal extent of significant decoding results as derived from a dependent-samples t-test (two-sided, p = 0.002, cluster corrected across time).

Memory performance was tested both before and after the sleep period in a step-wise manner. First, participants made word-recognition judgments (old or new). Then, for recognized words only, recall of the associated image exemplar (object or scene, depending on experimental session) was assessed. The resulting recall performance was then normalized by the amount of correctly recognized items (i.e. ‘hits’). To avoid any impact of pre-sleep testing on our behavioral consolidation measures ^41,42^, only half of the learned material was tested before sleep, while the remaining half was tested after sleep. Finally, at the end of the experimental sessions participants performed an independent ‘localizer task’, where a new set of object and scene images was presented (including both stimulus categories, irrespective of experimental session). This localizer served to train a linear classifier to distinguish object- vs. scene-related EEG patterns.

### Behavioral results & category classification during the localizer task

First, we calculated d-prime (d’ ^43^) as a general measure of recognition memory performance (for a detailed overview of memory measures as well as sleep characteristics see Supplementary Tables 1 and 2). Both pre- and post sleep d’ levels confirmed that participants could reliably discriminate between old and new items (i.e.: d’ > 0; pre-sleep objects: d’ = 2.11 ± 0.14, scenes: d’ = 2.02 ± 0.22; post-sleep objects: d’ = 1.76 ± 0.19, scenes: d’ = 1.69 ± 0.23). Out of hits, participants recalled the correct image for 64.31 ± 3.23% before sleep (objects: 64.90 ± 3.99%, scenes: 63.72 ± 5.20%) and for 57.61 ± 3.91% after sleep (objects: 59.39 ± 5.71%, scenes: 55.82 ± 5.47%).

To test for potential differences in memory performance between test times and stimulus categories, we conducted ANOVAs for recognition memory (d’) and cued recall, including the factors category (object vs. scene) and test-time (pre- vs. post-sleep). Results indicated that memory performance (both recognition and recall) declined over the course of sleep (main factor test-time: recognition memory: *F*_1,19_ = 10.91; p = 0.004; cued recall: *F*_1,19_ = 15.53; p = 0.001). Importantly though, no difference in memory performance between categories was observable (main effect category: recognition memory: *F*_1,19_ = 0.21; p = 0.65; cued recall: *F*_1,19_ = 0.38; p = 0.54) and no interaction between test-time and learning category (recognition memory: *F*_1,19_ = 0.003; p = 0.95; associative memory: *F*_1,19_ = 0.69; p = 0.41), ensuring that task difficulty was highly comparable between image categories (also see Table S1).

The localizer task at the end of each session was employed to derive the neural signatures of object vs. scene processing, which were then used to track category-specific memory reactivation during NREM sleep (see below). Participants were presented with novel sets of object and scene images and performed a continuous recognition task on these images. Specifically, each image was presented twice (mean distance between successive presentations = 8.06, range = 2 − 33) and participants were instructed to indicate whether a given item was ‘new’ (first presentation) or ‘old’ (second presentation). As expected, participants showed high accuracy levels on this task (objects: 97.02 ± 0.61 correct decisions; scenes: 92.57 ± 4.44 correct decisions), with performance again matched between image categories (t_(19)_ = 1.05, p = 0.31).

To extract the category-specific (i.e., object and scene) patterns of neuronal activity, we pooled the localizer data across experimental sessions and performed multivariate classification (linear discriminant analysis; LDA) on these data (Fig. 1c). Using 5-fold cross-validation (see Methods), above-chance classification accuracy emerged around 150 ms following image onset, was sustained until 2800 ms and peaked at 600 ms (p = 0.002, corrected for multiple comparisons across time). Hence, the localizer data allowed us to isolate brain patterns associated with the processing of object and scene images, which we then used to guide analysis of category-specific reactivation during sleep (for results concerning the stability of the decoding approach see Supplementary Fig. 1).

### Endogenous memory reactivation during NREM sleep is clocked by SO-spindle complexes

As mentioned above, theoretical models and recent empirical findings point to particular role of SO-spindle coupling for memory consolidation. We thus tested the resulting prediction that the joint presence of SOs and sleep spindles (henceforth referred to as ‘SO-spindle complexes’) would drive endogenous memory reactivation during human sleep. SOs and sleep spindles were detected in the EEG data using established algorithms ^8,44^. To isolate SO-spindle complexes, we identified events where SO down-states were followed by sleep spindles within a time window of 1.5 seconds (for a time-frequency representation of the SO-spindle complexes see Fig. 2a; for a peri-event SO-spindle histogram, see Supplementary Fig. 2). To determine whether learning-related (i.e. category-specific) neuronal activity would be differentially reactivated during SO-spindle complexes, we first trained a classifier on the concatenated localizer data from both experimental sessions [−0.5 to 3 seconds]. Importantly, the localizer tasks of both sessions included object and scene images, to ensure that multivariate measures of potential reactivation not merely reflect session-specific EEG properties. The resulting training weights were then applied on both sessions’ sleep data, centered around the down-state of SO-spindle complexes (for related results were the data was locked to different spindle features see Supplementary Fig. 3). Classifier testing labels reflected the stimulus category used in the preceding encoding session (object or scene), such that above-chance classification signifies endogenous activation patterns more strongly resembling the just-learned stimulus category than the alternative stimulus category.

As shown in Fig. 2b, results revealed a cluster of significant above-chance classification from 800 to 1200 ms relative to the SO down-state (p = 0.016, corrected for multiple comparisons across time, localizer time-window [1000 to 1800ms]), emerging between maximum and offset of coupled sleep spindles (for the corresponding accuracy map see Supplementary Fig. 4; for participant specific classification values see Supplementary Table 3). No negative cluster survived correction for multiple comparisons (cluster with smallest p >0.6).

**Figure 2.**
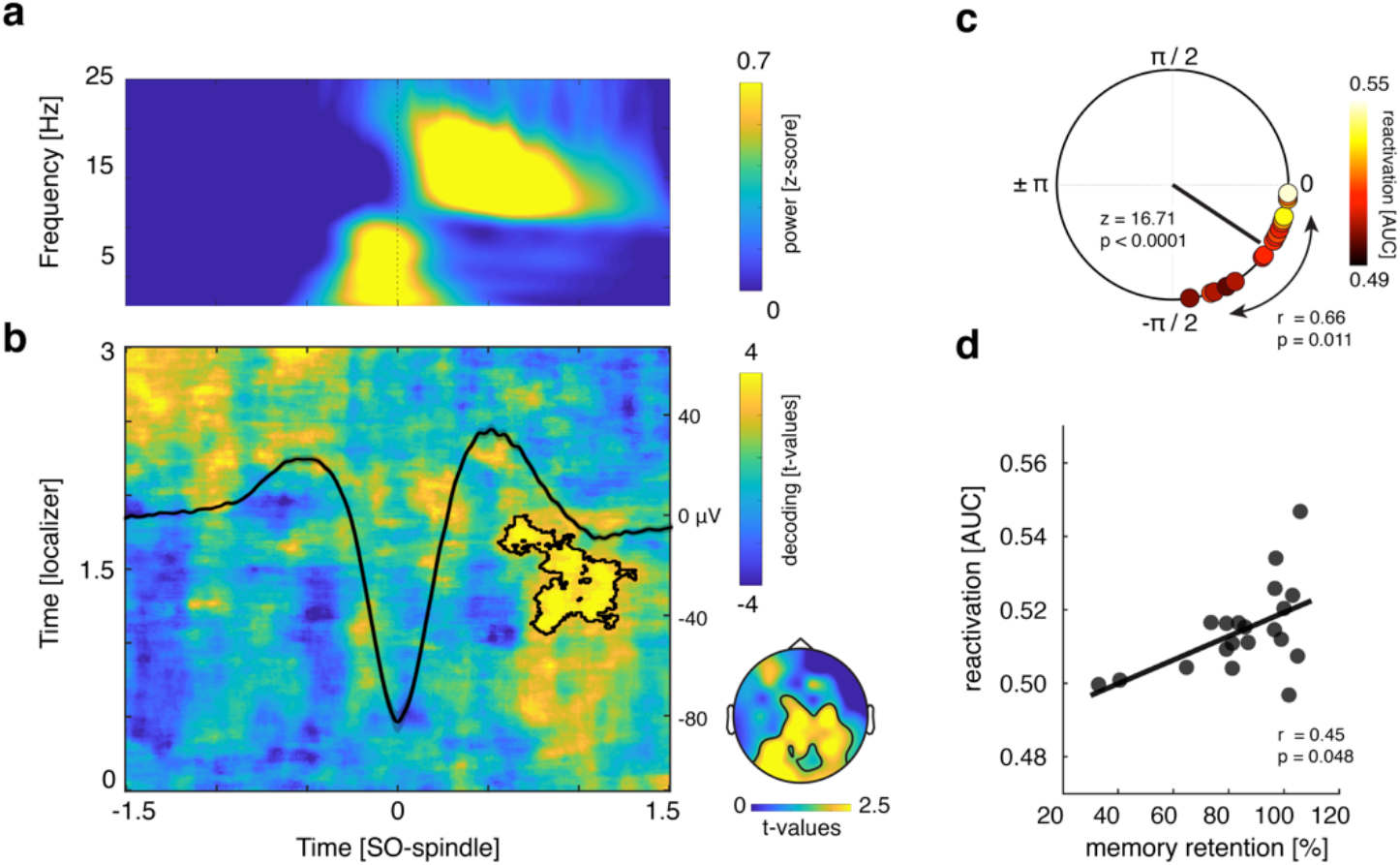
SO-spindle locked memory reactivation. (a) Time frequency representation of all SO-spindle segments (z-scored across time; only positive values are displayed, with yellow indicating power increases). (b) Learning-related brain patterns (objects vs. scenes) were decodable during SO-spindle complexes (contour lines indicate the extent of the significant cluster, p = 0.016 corrected; color range (blue to yellow) represents t-values against surrogate decoding performance, which was estimated by shuffling the training labels 250 times). The averaged EEG trace (all instances in which SO down-states were followed by sleep spindles within 1.5 seconds at channel Cz in microvolt [μV]) illustrates the relationship of the observed reactivation signal with ongoing oscillatory activity. The topographical insert illustrates the results of a ‘searchlight decoding procedure’, indicating that bilateral parietal and occipital areas exhibited stimulus-category related effects (please note that statistical tests were done for illustrative purposes only). (c) Phases of the SO-spindle modulation derived from channel Cz, illustrating the clustering of spindle power toward the SO up-state (up-state corresponding to 0 and down-state to ± π, with –π/2 reflecting the down-to up-state transition; Rayleigh test: p < 0.0001; z = 16.71). The black line illustrates the mean coupling direction and vector length (−36.78° ± 5.48°, mean vector length = 0.91). Circular-linear correlation analysis between the individual mean SO-spindle coupling phase (circles) and the mean reactivation strength (area under the curve [AUC] scores; color coded, with white indicating high classification performance and black low classification performance) revealed a positive association (r = 0.66; p = 0.011). (d) Reactivation strength correlated positively with behavioral levels of associative memory consolidation (Spearman’s Rank Correlation, r = 0.45, p = 0.048).

But does endogenous memory reactivation indeed requires the joint presence of SOs and spindles? To address this question, we performed the same decoding procedure, but locking the data to solitary SO or spindle events (thus, SOs without spindles and vice versa). For both types of events, when testing accuracy levels against chance at any localizer time x sleep time point, no significant cluster of above chance classification emerged (in both cases cluster with the smallest p > 0.2, see Supplementary Fig. 5; similarly, testing the classifier on Slow spindle – SO locked data did not yield any significant cluster of above-chance classification (cluster with smallest p = 0.67; see Supplementary Fig. 6)).

### Precision of SO-spindle coupling correlates with reactivation strength

If SO-spindle coupling is indeed instrumental for consolidation, its precision should impact the extent of endogenous memory reactivation. To quantify the preferred phase of SO-spindle modulation, we determined in every participant the SO phases corresponding to the spindle peak amplitudes (electrode Cz). In 16/20 participants we found significant non-uniform distributions (p < 0.05; Rayleigh test, mean vector length: 0.34 ± 0.03). In line with previous findings, we found a significant non-uniform distribution across participants (Rayleigh z = 16.71, p < 0.0001), with spindles peaking near the SO up-state (corresponding to 0°; mean coupling direction: −36.78° ± 5.48°; see Figure 2c).

To further test whether the precision of SO-spindle coupling would be relevant for the reactivation of memories we computed a circular-linear correlation between each participant’s preferred SO-spindle phase (averaged across sessions) and their mean reactivation strength (averaged across the significant cluster shown in Fig. 2b). The individual SO-spindle modulation phase was significantly correlated with decoding accuracy (r = 0.66; p = 0.011). The distribution indicated that the closer the spindles were nested towards the SO-upstate, the higher the fidelity of the associated reactivation signal (see Figure 2c, for a scatter plot see Supplementary Fig. 7; for additional analyses estimating the impact of trait-like characteristics in this context, see Supplementary Results). To ensure that the results described above were not driven by differential wake classification characteristics, we conducted a partial circular-linear correlation with the mean decoding levels from the localizer tasks (averaged across the significant cluster shown in Fig. 1c) as a covariate. Again, we observed a positive relationship between the individual SO-spindle modulation phase and decoding accuracy (r = 0.65; p = 0.012).

### Reactivation strength predicts consolidation of associative memories

If SO-spindle triggered reactivation reflects memory-related processes, one would expect a functional link with behavioral expressions of consolidation. To address this question, we correlated, across participants, levels of post-sleep memory retention and reactivation strength. Specifically, a ‘retention index’ (proportion of post-sleep recalled images (out of hits) in relation to pre-sleep memory performance; see Methods section for details) was collapsed across sessions and correlated with decoding accuracies averaged across the significant cluster reported above. As shown in Figure 2d, we observed a significant positive relationship between the two variables (Spearman rho = 0.45, p = 0.048). Of note, no association between decoding accuracy and recognition memory performance was detectable (r = 0.02, p = 0.93), indicating that reactivation strength was specifically linked to the consolidation of hippocampal-dependent associative memories ^45^. However, the correlation between reactivation and consolidation of associative memory was not significantly greater than that with recognition memory (z = 1.35; p = 0.17). Lastly, we again controlled this analysis for localizer decoding levels using a partial correlation, which substantiated the results (Spearman rho = 0.45, p = 0.049).

## DISCUSSION

Our results demonstrate that consolidation relies on endogenous memory reactivation clocked by SO-spindle complexes. In particular, we found that during the presence of SO-spindle complexes, activation patterns were biased towards the previously encoded learning material (Fig 2a-b). Moreover, the precision of SO-spindle coupling predicted the fidelity of memory reactivation (Fig 2c). Finally, reactivation strength predicted the amount of consolidation across participants, highlighting its functional significance for behavior (Fig 3d).

NREM sleep oscillations (SOs, spindles and ripples) have long been implicated in the memory function of sleep, and recent work has emphasized the importance of their temporal synchronization ^46^. Specifically, the precise timing of SOs, spindles and ripples is thought to enable the relay of hippocampus-dependent memories to cortical networks ^1^. Indeed, recent work in rodents has shown that their co-occurrence is necessary for effective consolidation as assessed via fear conditioning ^10^ or an object-in-place recognition task ^9^. However, how these tasks relate to expressions of episodic memory in humans is not entirely clear. Human iEEG work with epilepsy patients has corroborated the triple-interaction of these sleep oscillations ^8,24,27^, but none of these studies has assessed memory reactivation or the effects on behavior. Investigation of healthy participants via scalp EEG has shown that brain patterns across sleep differ as a function of prior learning tasks ^47^, but these activation patterns were not directly related to wake activity or to discrete SOs/spindles. Another study employed simultaneous EEG-fMRI and found univariate signal increases in learning-related areas during spindles ^48^ (see also ^44^), but it remained open whether such reactivation bears relevance for memory consolidation. Finally, the advent of targeted memory reactivation (TMR) protocols ^49,50^ has shown evidence for both SO-spindle complexes and information processing in response to external reminders ^36,38,51–55^, but it is unclear whether and how such exogenous memory reactivation relates to endogenous reactivation in service of memory consolidation. In sum, different lines of research across species point to a key role in coupled sleep oscillations, but the dynamics of endogenous reactivation in humans and its relevance for memory consolidation has remained unclear.

In the current study, we tackled this question by employing two learning sessions per participant, each using different and analytically discriminable learning stimuli (object and scene images, Fig.1a). To ensure that multivariate measures of reactivation not merely reflect session-specific EEG properties, we included an object/scene localizer task in each session and trained a linear classifier on the combined data. This allowed us to track the re-emergence of learning categories during the nap periods. It deserves mention that decoding levels were modest in general and not every participant reached above-chance classification (18/20, see Figure 2d, Supplementary Table 3). Several reasons might limit the effect size when decoding memory reprocessing during sleep. First, the signal of interest (i.e. sleep electrophysiology) is inherently noisy. Guided by theoretical considerations we limited the search-space for memory reactivation to the presence of SO-spindle complexes. Still, it is unlikely that each single SO-spindle complex is associated with memory reactivation. Including the presence of ripples as a criterion may increase sensitivity, but even SO-spindle-ripple complexes are unlikely to yield robust memory reactivation in every instance ^56^. Second, our data show that SO down-states represent viable reference points for time-locking the analysis of memory reactivation. However, there is considerable variability in signal characteristics across SOs and spindles (e.g. event durations or peak times), and such across-event variability diminishes classification power which relies on spatiotemporal activation patterns common across events. That said, decoding levels observed here are in line with previous TMR studies examining sleep-related memory reactivation with multivariate classification ^36,53,57^. Importantly, we found that higher decoding performance correlates with the behavioural expression of memory consolidation across participants, further corroborating the functional significance of reactivation.

Another key feature of our paradigm was the assessment of both item- and associative memory performance. Interestingly, the strength of memory reactivation during sleep predicted consolidation levels for associative memory only. This finding could indicate that reactivation particularly benefits hippocampus-dependent memories ^45^. However, it might also reflect the fact that reactivation pertained to the categorical features of the learning material, which was also the aspect relevant for associative- and not item memory. Moreover, while performance levels were carefully matched between object and scene tasks (Fig. 1b), performance was lower for associative memory than for item recognition. Thus, differential effects of reactivation for associative- vs. item memory could also suggest differential benefits of sleep for weaker vs. stronger memories ^58–61^ but see ^62^.

Owing to the limited spatial resolution of scalp EEG (especially for transient high-frequency oscillations), our current data remain agnostic with regard to hippocampal ripples. That said, a recent iEEG study has shown that both hippocampal ripples and hippocampal-cortical interactions are most eminent when preceded by a cortical SO-spindle complex ^24^. To the extent that reactivation observed here is linked to hippocampal engagement, the timing of our effects (Fig.2 a-b) is consistent with accumulating evidence that the hippocampal-cortical dialogue is in fact initiated by cortex ^24,25,63–65^. One tentative interpretation of our results might thus be that cortical SO-spindle complexes trigger hippocampal memory reactivation while ensuring that the cortical target area is optimally tuned for synaptic plasticity and memory reprocessing 19,21,66. Indeed, recent rodent work has shown that optogenetic induction of SO-locked spindles enhances SOs-spindle-ripple coupling and the consolidation of hippocampus-dependent memories ^10^. Our finding that reactivation peaks towards the end of spindles (Figure 2b) is consistent with the idea that mnemonic reprocessing and integration into neocortical networks continue after sleep spindles, i.e. during periods of spindle ‘refractoriness’ ^67^. Likewise, intracranial recordings in humans have shown that hippocampal-cortical connectivity (‘mutual information’) mediated by hippocampal ripples occurred ~500-1500 ms after the SO down-state ^24^, again matching the time window in which we observed memory reactivation. Together, one tentative scenario might be that memory processing is most beneficial after SO-spindle complexes, i.e. at time points of elevated cortical plasticity.

Analytically, our approach relied on (i) matching behavioural performance between sessions, (ii) pooling sleep data across both sessions and (iii) deriving evidence for the reactivation of learning material across all aggregated SO-spindle complexes. These design features leave some interesting questions open for future work. First, to what extent might trait-like participant characteristics drive both reactivation and memory processes? Using our sleep questionnaires, we were able to rule out subjective sleep quality and circadian rhythm as confounds (see Supplementary Results), but there may be other trait-like factors impacting reactivation and consolidation. An alternative design would be to conduct a longitudinal study in which within-participant levels of learning and consolidation are experimentally manipulated across multiple sessions (e.g. by varying encoding depth or task difficulty). Second, while aggregating all SO-spindle events is essential for the classification approach, it leaves open whether reactivation occurs during each SO-spindle event. An alternative approach might be to use intracranial recordings to identify single neurons that are tuned to stimuli used in a specific learning session and then track engagement of these neurons during individual SO-spindle complexes. Such more fine-grained methods might provide additional insights into reactivation-related characteristics (e.g. accuracy and frequency of reactivation processes). In conclusion, our results indicate that endogenous memory reactivation in service of sleep-dependent consolidation is clocked by the fine-tuned coupling of SOs and spindles. Future work employing simultaneous recordings from the hippocampus will further elucidate the intricate dynamics underlying the hippocampal-cortical dialogue of systems consolidation.

## METHODS

### Participants

Twenty healthy, right-handed participants (mean age: 20.75 ± 0.35; 17 female) with normal or corrected-to-normal vision took part in the experiment. An additional five participants had to be excluded due to insufficient sleep (less than 30 minutes sleep during one of the sessions). The sample size was determined in accordance with previous human sleep and memory studies (e.g. ^30,68^). Pre-study screening questionnaires (including the Pittsburgh Sleep Quality Index (PSQI, ^69^), the Morningness-Eveningness Questionnaire ^70^ and a self-developed questionnaire querying general health status and the use of stimulants) indicated that participants did not take any medication at the time of the experimental session and did not suffer from any neurological or psychiatric disorders. All participants reported good overall sleep quality. Furthermore, they had not been on a night shift for at least 8 weeks before the experiment. All participants were instructed to wake up by 7:00 am and avoid alcohol the evening before and caffeine on the day of the experimental sessions. They confirmed at the beginning of each experimental session their adherence to the requirements. The study was approved by the University of Birmingham Research Ethics Committee and written informed consent was obtained from participants.

### Stimuli and procedures

#### Overview

The experiment consisted of two experimental sessions (object and scene condition), separated by at least one week (mean = 8.5 ± 0.85 days). The order of the two sessions was counterbalanced across participants. On experimental days participants arrived at the sleep laboratory at 11am. The experimental session started with the set-up for polysomnographic recordings during which electrodes for electroencephalographic (EEG), electromyographic (EMG) and electrocardiographic (ECG) recordings were applied. Before the experimental sessions, participants were habituated to the environment by spending an adaptation nap in the sleep laboratory. At around 12:00 am the experiment started with a modified version of the psychomotor vigilance task (“PVT” ^71^), followed by the memory task (for details see Memory Task below). The sleep period began at ~1pm and participants were given 120 minutes to nap (mean total sleep time: 101.63 ± 2.23 minutes; for sleep characteristics see Supplementary Table 2). Afterwards, the vigilance of all participants was assessed using the PVT and memory performance was tested again. At the end of each session a localizer task was conducted (see Localizer Task for details).

#### Stimuli

A set of in total 360 verbs and 240 images (half objects, half scenes) served as experimental stimuli during both sessions. Objects were images of animals, food, clothing, tools or household items presented on a plain white background (e.g. a hammer). Scenes were images of nameable landscapes or places (e.g., a coffee shop). All images were taken from ^72^.

#### Experimental Tasks

For the recording of behavioral responses and the presentation of all experimental tasks, Psychophysics Toolbox Version 3 ^73^ and MATLAB 2018b (MathWorks, Natick, USA) were used. Participants completed a practice run (5 trials) of each experimental task in advance to ensure they fully understood the instructions. Responses were made via keyboard presses on a dedicated PC. Across all experimental phases, presentation order of stimuli was randomized across participants.

#### Psychomotor Vigilance Task

The vigilance of the participants was assessed using a modified version of the psychomotor vigilance task (“PVT” ^71^) before the encoding phase and right after the sleep period. Participants were presented with a centred fixation cross on the computer screen. Every 2-10 s the fixation cross was replaced by a counter counting up from 0 to 2 s in steps of 20 ms. Participants were instructed to stop the counter as fast as possible by pressing the space bar. After each trial participants were provided with feedback about their reaction time. The task was administered for 5 minutes. For PVT related results see Supplementary Fig. 8.

#### Familiarization

The experiment began with an image familiarization phase. The purpose of this part was (i) to facilitate learning of the verb-image pairs in the main encoding session and (ii) to provide the proper image names for subsequent cued recall. Each trial started with a fixation cross, presented for 1.5 ± 0.1 seconds. Subsequently, participants saw one of 130 images showing objects or scenes (depending on the experimental session). 120 of these images were part of the subsequent learning material and were accompanied by a caption naming the exemplar. 10 additional images, which were not further used during the experiment, were accompanied by an erroneous description. Each stimulus combination was presented for 2.5 seconds on the computer screen. The participants’ task was to press a button whenever they encountered a wrong image-word combination.

#### Encoding

Participants learned pairwise associations between 120 verbs and images. The images comprised either objects or scenes (depending on experimental session).

Each trial started with a fixation cross, presented for 1.5 ± 0.1 seconds. Afterwards, a verb (e.g., ‘‘jump’’) was presented for 1 second on the computer screen and immediately followed by the to-be-associated image for 4 seconds. Participants were instructed to form a vivid mental image or story linking the verb and the object/scene. After the presentation of the image (4 seconds), they had to indicate whether the image they had formed was realistic or bizarre. In addition, participants were informed that their memory performance for verb- image pairs would be tested later. The learning block was run twice with varying trial order to reach satisfactory levels of pre-sleep memory performance (as determined in a pilot study).

#### Pre-sleep Memory Test

In order to prevent any testing effect on our behavioral measures of memory consolidation ^41,42^, only half of the learned verb-image combinations was tested during the pre-sleep memory test. Thus, the pre-sleep memory test included 60 randomly chosen verbs intermixed with 30 new verbs, which were not seen by the participants before (‘foils’). Each trial started with a fixation cross, presented for 1.5 ± 0.1 seconds. After the fixation cross, a verb was presented on the computer screen. After 3 seconds, participants had to indicate whether the verb was ‘old’ (i.e., part of the learning material) or ‘new’ (i.e., it was not seen during learning) within the next 10 seconds. In case of ‘new’ responses, participants immediately moved on to the next trial. In case of ‘‘old’’ responses, participants were required to type a description of the image they had in mind or to type ‘do not know’ in case they could not recall the target image. Trials were coded as correct if (i) the participant typed the same caption as shown during the familiarization phase or (ii) the description unambiguously matched the content of the image

#### Sleep period

The nap period began at ~1pm. Participants had the opportunity to sleep in a laboratory bedroom for 120 minutes, while their brain activity was monitored using polysomnography).

#### Post-sleep Memory Test

20 minutes after waking up, participants performed another memory test on the remaining 60 study items. This followed the same procedures as the pre-sleep memory test with the exception that new foil verbs were used.

#### Localizer Task

During the localizer task participants were presented with a new set of images comprising objects and scenes (90 objects and 90 scenes, irrespective of session). Each trial started with a fixation cross, presented for 1.5 ± 0.1 seconds. Subsequently, a randomly chosen image (object or scene) was presented on the computer screen for a minimum of 2.5 and a maximum of 10 seconds. Each image was presented twice during the task and participants were instructed to indicate whether it was shown for the first (‘new’) or second (‘old’) time (mean distance between successive presentations = 8.06, range = 2 − 33).

By administering the localizer task at the very end of each session, we assured that participants engaged exclusively with a given stimulus category before sleep (objects or scenes, respectively). The rationale of this approach was to keep the category-specific representations during learning as pure as possible, in an effort to bias their reactivation during the subsequent sleep period. However, presenting both stimulus categories during the localizer task ensured that category-specific classifier evidence during sleep would not merely reflect general differences between sessions (e.g. electrode impedances, electrode positions etc.).

### EEG

A Brain Products 64 channel EEG system was used to record electroencephalography (EEG) throughout the experiment. Impedances were kept below 10 kΩ. EEG signals were referenced online to electrode FCz and sampled at a rate of 1000 Hz. Furthermore, EMG and the ECG was recorded for polysomnography. Sleep architecture was determined offline according to standard criteria by two independent raters ^74^.

### Data analysis

#### Behavioral preprocessing

To assess recognition memory performance, we calculated the sensitivity index d’ [i.e., z(Hits) – z(False Alarms)] according to signal detection theory. Proportions of 0 and 1 were replaced by 1/2N and 1-1/2N, respectively, with N representing the number of trials in each proportion (i.e., N = 60, see ^43^).

For associative memory performance we calculated the proportion of correctly recalled images relative to the number of recognized words (i.e. (recalled images / hits)*100). To correlate levels of memory retention and reactivation strength we derived a ‘retention index’. We computed the proportion of post-sleep recalled images (out of hits) in relation to pre-sleep memory performance (i.e. (recalled out of hits post-sleep / recalled out of hits pre-sleep)*100) and collapsed these measures across sessions.

#### EEG Data Analysis

EEG data were preprocessed using the FieldTrip toolbox for EEG/MEG analysis ^75^. All data were downsampled to 200 Hz. Subsequently, the localizer and sleep data were segmented into epochs. The temporal range of the epochs was [−1 to 3] seconds around stimulus onset for localizer trials. As in other studies concentrating on the coordination of SOs and spindles (e.g ^8,30–32,76^) we specifically focused on electrode Cz due to the spatial distribution of both oscillations. Both oscillations show strong presence over central areas, rendering Cz an optimal target zone for investigating concomitant activity of SOs and (fast) spindles. Hence, for the sleep data, slow oscillation – spindle epochs [−2.5 to +2.5 seconds] time-locked to SO down-states were extracted from channel Cz (for details see Event detection).

Noisy EEG channels were identified by visual inspection, discarded and interpolated, using a weighted average of the neighboring channels. The localizer data were additionally subjected to an independent component analysis ^77^ and ICA components associated with eye blinks and eye movements were identified and rejected.

#### Event detection and SO-spindle coupling

SOs and sleep spindles were identified for each participant, based on established detection algorithms ^8,44^. Following standard procedures, all sleep data were re-referenced against linked mastoids for sleep scoring and event detection ^74,78,79^; please note that the classification results reported in Fig 2b remained unchanged when using a common average reference scheme). SOs were detected as follows: Data were filtered between 0.3–1.25 Hz (two-pass FIR bandpass filter, order = 3 cycles of the low frequency cut-off). Only movement-free data (as determined during sleep scoring) from NREM sleep stages 2 and 3 were taken into account. All zero-crossings were determined in the filtered signal at channel Cz, and event duration was determined for SO candidates (that is, down-states followed by up-states) as time between two successive positive- to-negative zero-crossings. Events that met the SO duration criteria (minimum of 0.8 and maximum of 2 s, 0.5–1.25 Hz) entered the analysis. 5-s-long segments (±2.5 s centered on the down-state) were extracted from the unfiltered raw signal.

For spindle detection, data were filtered between 12–18 Hz ^25,80^ (two-pass FIR bandpass filter, order = 3 cycles of the low frequency cut-off), and again only artifact-free data from NREM sleep stages 2 and 3 were used for event detection. The root-mean-square (RMS) signal was calculated for the filtered signal at channel Cz using a moving average of 200 ms, and a spindle amplitude criterion was defined as the 75% percentile of RMS values. Whenever the signal exceeded this threshold for more than 0.5 s but less than 3 s (duration criteria), a spindle event was detected. Epochs time-locked to the minimum spindle trough (−2.5 to +2.5 seconds) were extracted from the unfiltered raw signal for all events. To isolate SO-spindle complexes, we determined for all SOs whether a spindle was detected following the SO (SO down-state + 1.5 seconds). Finally, SO-spindle events were extracted (−2.5 to +2.5 seconds with regards to the SO-down-state) from the raw signal at channel Cz.

For the analysis of SO-spindle coupling ^8,24^, we filtered the SO-spindle data in the SO range (0.3 − 1.25 Hz, two-pass Butterworth bandpass filter), applied a Hilbert transform and extracted the instantaneous phase angle. Next we filtered the same data segments in the spindle range (12-18 Hz two-pass Butterworth bandpass filter), Hilbert transformed the signal and extracted the instantaneous amplitude. Only data points within ±1.5 seconds were considered to avoid filter-related edge artifacts. Then we detected the maximal sleep spindle amplitude in channel Cz and isolated the corresponding SO phase angle. The preferred phase of SO-spindle coupling was then obtained from averaging all individual events’ preferred phases of each participant, and the resulting distribution across participants was tested against uniformity (Rayleigh test, CircStat toolbox ^81^).

#### Multivariate analysis

Multivariate classification of single-trial EEG data was performed using MVPA-Light, a MATLAB-based toolbox for multivariate pattern analysis ^82^. For all multivariate analyses, a linear discriminant analysis (LDA) was used as a classifier ^82^. Prior to classification, all data were re-referenced using a Common Average Reference (CAR).

##### Classification within the localizer task

The localizer data were z-scored across all trials for each time point separately. Next, data from both sessions were collapsed and subjected to a principal component analysis (PCA), which transforms the data into linearly uncorrelated components, ordered by the amount of variance explained by each component ^83^. PCA was applied to reduce dimensionality and limit over-fitting ^84^ and the first 30 principal components were retained for further analysis ^85–87^. To quantify whether object and scene representations can be differentiated in the localizer, the classifier was trained and tested to discriminate between object and scene trials. Data were smoothed using a running average window of 150 ms. The EEG channels served as features and a different classifier was trained and tested on every time point. As metric, we used Area Under the ROC Curve (AUC), which indexes the mean accuracy with which a randomly chosen pair of Class A and Class B trials could be assigned to their correct classes (0.5 = random performance; 1.0 = perfect performance). To avoid overfitting, data were split into training and test sets using 5-fold cross-validation ^88^. Since cross-validation results are stochastic due to the random assignment of trials into folds, the analysis was repeated five times and results were averaged. For statistical evaluation, surrogate decoding performance was calculated by shuffling the training labels 250 times. Resulting surrogate performance values were then averaged, providing baseline values for each participant under the null hypothesis of label exchangeability

##### Decoding object and scene representations during SO-spindle complexes

To investigate differential evidence for object vs. scene representations as a function of prior learning during SO-spindle complexes (Figure 2b), we used the temporal generalization method ^89^. Prior to decoding, a baseline correction was applied based on the whole trial ([−0.5 to 3 seconds] for localizer segments; [−1.5 to 1.5 seconds] for SO-spindle segments). Next, localizer and sleep data were z-scored across trials and collapsed across sessions. PCA was applied to the pooled wake-sleep data and the first 30 principal components were retained. Localizer and sleep data were smoothed using a running average window of 150 ms. A classifier was then trained for every time point in the localizer data (Figure 2b, vertical axis) and applied on every time point during SO-spindle complexes (horizontal axis). No cross-validation was required since localizer and sleep datasets were independent. As metric, we again used AUC (see above). For statistical evaluation, surrogate decoding performance was calculated by shuffling the training labels (stemming from the localizer task) 250 times. Again, the resulting performance values were averaged, providing baseline values for each participant under the null hypothesis of label exchangeability

##### Searchlight analysis

To resolve the topography of diagnostic features, we conducted a ‘searchlight decoding procedure’. In brief, PCA components were projected back to sensor space and the classification procedure was repeated across moving kernels of small electrode clusters, with neighboring electrodes being selected as features [feature number range: 5 to 9]. Classifiers were trained for every time-point in the localizer data and applied on every time point during SO-spindle complexes. Finally, classification values were collapsed across our time windows of interest [localizer time: 1000 to 2000 ms; SO-spindle time: 800 to 1200 ms] and tested against chance level (corrected for multiple comparisons across space). A broad cluster of above-chance classification comprising bilateral parietal and occipital areas emerged (p_cluster_ = 0.004).

#### Time-frequency analysis

Time-frequency analysis of the SO-spindle segments was performed using FieldTrip. Frequency decomposition of the data, using Fourier analysis based on sliding time windows (moving forward in 50-ms increments). The window length was set to 5 cycles of a given frequency (frequency range: 1 – 30Hz in 1Hz steps). The windowed data segments were multiplied with a Hanning taper before Fourier analysis. Afterwards, power values were z-scored across time [−4 to 4 seconds]. The longer time segments were chosen to allow for resolving low frequency activity within the time-windows of interest [−1.5 to 1.5 seconds] and avoid edge artifacts.

### Statistics

Behavioral retrieval data were subjected to a 2 (Category: Object/Scene) X 2 (Test-Time: Pre-sleep/Post-sleep) repeated measures ANOVA. To test for potential differences in memory accuracy between sessions in the localizer task, a paired sampled t-test was computed. The statistical significance thresholds for all behavioral analyses were set at p <. .05. Spearman correlation was used to assess the relationship between memory retention and reactivation strength. To control for mean decoding levels from the localizer tasks (averaged across the significant cluster), a partial Spearman correlation was used. SPSS (IBM Corp., Version 26) and Matlab was used for behavioral data analyses.

FieldTrip’s cluster permutation test ^90^ was used to deal with the multiple comparisons problem for all classification analyses. A dependent-samples t-test was used at the sample level to identify clusters of contiguous time-points across participants and values were thresholded at p = .05. Maxsum (sum of all t-values in cluster) served as cluster statistic and Monte Carlo simulations were used to calculate the cluster p-value (alpha = .05, two-tailed) under the permutation distribution. Analyses were performed at the group level. The input data were either classification values across time (Figure 1c) or time x time classification values (Figures 2b). In all cases a two-sided cluster permutation test with 1000 randomizations was used to contrast classification accuracy against chance performance.

Non-uniformity of the preferred phase with regard to SO-spindle coupling was assessed using the Rayleigh test (CircStat toolbox). The non-linear relationship between SO-spindle coupling and reactivation strength was determined with a circular linear correlation as implemented in the CircStat toolbox. A partial circular linear correlation modified from the CircStat toolbox was used to control for the mean decoding levels from the localizer task. In all cases the statistical significance thresholds were set at p < .05.

## Data availability

All relevant data supporting the main findings of this study are publicly available via the Open Science Framework (OSF: https://osf.io/gy25u). Source data are provided with this paper. A reporting summary for this article is available as a Supplementary Information file.

## Code availability

Code related to the main analyses of the manuscript is available at https://osf.io/gy25u/.

## Acknowledgments

This work was supported by a Wellcome Trust/Royal Society Sir Henry Dale Fellowship (107672/Z/15/Z) to B.P.S.. T.S. was supported by a grant of the Swiss National Science Foundation (SNSF) (P300P1_174450). T.St. research is funded by the European Research Council (https://erc.europa.eu/, Starting Grant 802681).

## Contributions

T.S., M.P. and B.S. conceived the study and designed the experiment. T.S. conducted the experiment. T.S., M.P., T.St. and B.S. analyzed the data and wrote the paper.

## Supplementary Information

**Supplementary Fig. 1:**
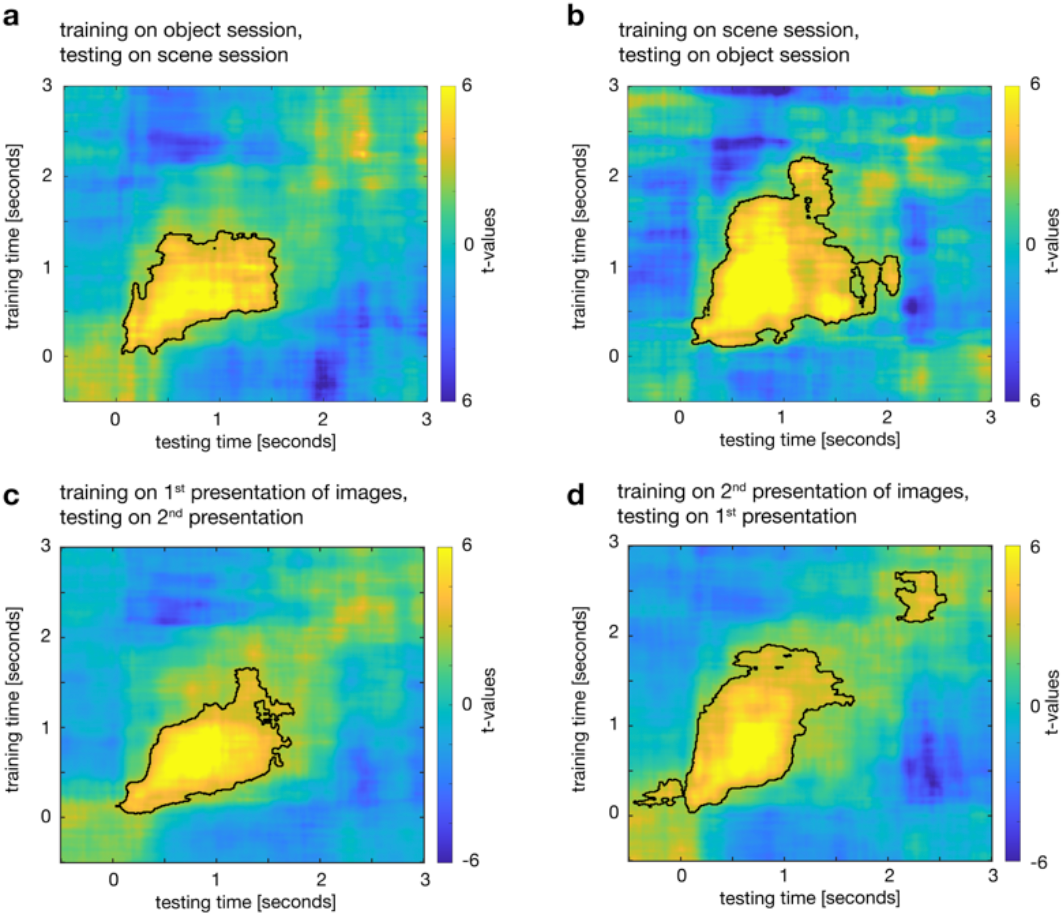
Stability of EEG object vs. scene decoding during the localizer tasks. (a + b) To estimate the across-session stability of our decoding approach, we trained a classifier on the localizer data of object sessions and tested it on the localizer data of the scene sessions. (a) We found robust above-chance classification when training the classifier on the localizer of the object session and applying the training weights to the corresponding data from the scene session (two-sided dependent-samples t-test; p_cluster_ = 0.002, cluster corrected across time). (b) The same result pattern emerged when training on the scene session data and applying the training weights to the object session data (two-sided dependent-samples t-test; p = 0.002, cluster corrected across time). (c + d) During the localizer task, each image was presented twice. To estimate the across-presentation stability of our decoding approach, we trained a classifier on the first image presentation tested it on the second presentation. (c) We observed a significant cluster of above-chance classification when training the classifier on the first presentation and applying it to the second presentation (two-sided dependent-samples t-test; p_cluster_ = 0.0018, cluster corrected across time). (d) A highly comparable result pattern emerged when training on the second presentation and applying it to the first presentation (two-sided dependent-samples t-test; p_cluster1_ = 0.002; p_cluster2_ =0.019, cluster corrected across time).

**Supplementary Fig 2.**
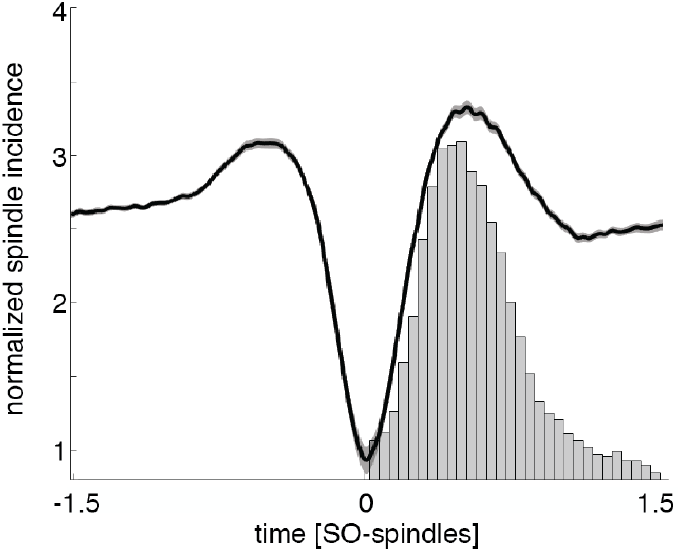
Perievent histogram of sleep spindles (amplitude maxima) following SO down-states (time = zero; normalized by number of spindles). Note that due to our definition of SO-spindle complexes (with sleep spindles following SOs), no spindles appear prior to SO down-states (time-point zero).

**Supplementary Fig. 3.**
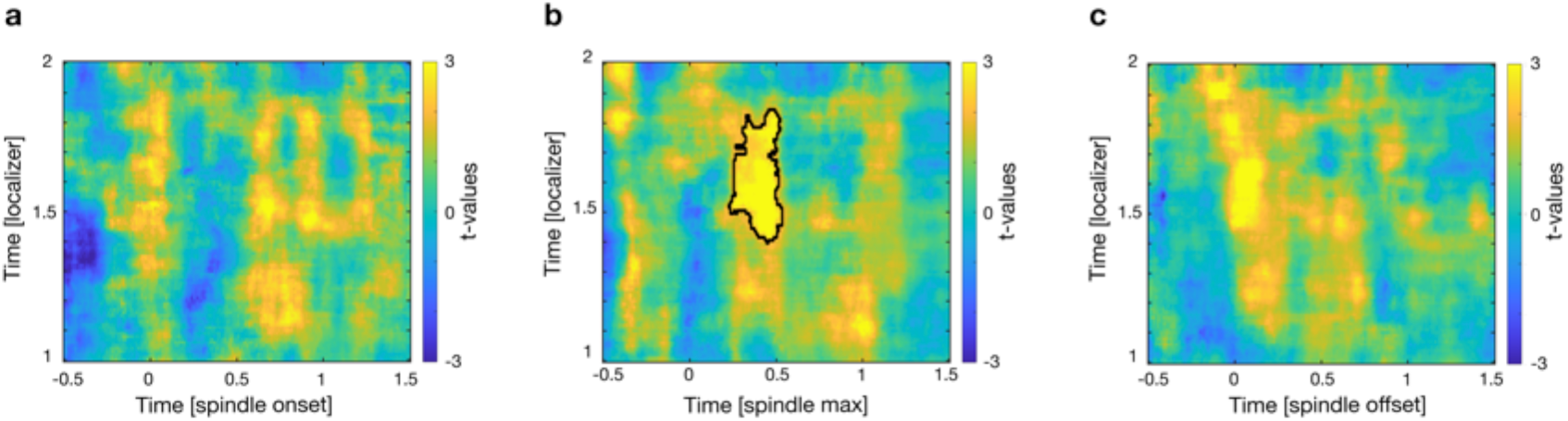
Classification locked to different spindle features. To test for the impact of different features of SO-spindle complexes on the synchronisation of reactivation events, the classification was computed on SO-spindle data locked to the onset, maximum and offset of spindles. Informed by the main analysis, the utilized time-window of both the localizer task and SO-spindle complexes were adjusted accordingly [localizer time: 1000-2000ms; SO-spindle time: −500 to 1500ms relative to respective feature (spindle onset, maximum amplitude and offset)]. Testing accuracy levels against chance at any localizer time x sleep time point for data locked to spindle on-and offsets (a + c) did not lead to any significant above chance classification (two-sided dependent-samples t-test; cluster with the smallest p-value for spindle onset: p = 0.25; spindle offset: p = 0.051, cluster corrected across time). However, testing the classifier on SO-spindle data locked to the spindle maximum (b) peaks yielded a positive cluster of significant above chance classification (two-sided dependent-samples t-test; p = 0.019, SO-spindle time [250 to 500ms], localizer time-window [1400 to 1800ms], cluster corrected across time).

**Supplementary Fig. 4.**
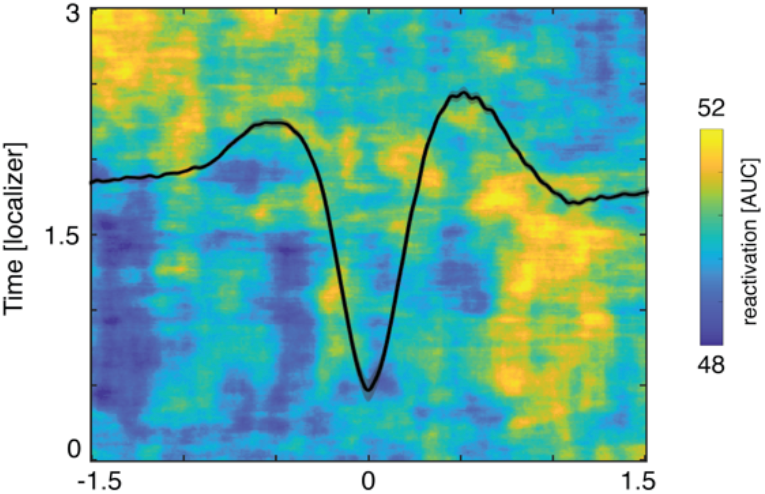
Accuracy map for the classification during SO-spindle complexes

**Supplementary Fig. 5.**
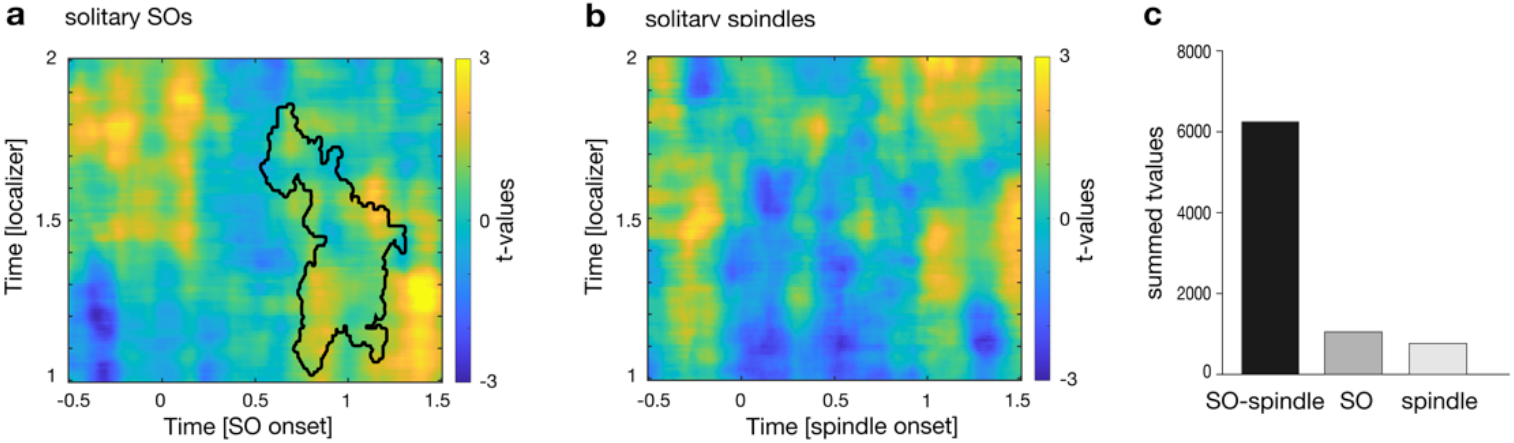
Memory reactivation during solitary SO and spindle events **(a + b)** To test whether endogenous memory reactivation indeed requires the joint presence of SOs and spindles, we performed the decoding procedure on solitary SO or spindle events (thus, SOs without spindles and vice versa). Time-window of both the localizer task and SO-spindle complexes were restricted according to the main results [localizer time: 1000-2000ms; SO-spindle time: −500 to 1500ms relative to respective event (SO down-state, spindle maximum)]. For both types of events, when testing accuracy levels against chance at any localizer time x sleep time point, no significant cluster of above chance classification emerged (two-sided dependent-samples t-test: in both cases cluster with the smallest p > 0.2; cluster corrected across time). The black contour lines in (a) illustrate the extent of the significant cluster derived from the main analysis (classification during the presence of SO-spindle complexes). (c) Summed t-values of the significant classification cluster (as derived from the main analysis, corresponding to the black contour lines in Fig 2b and Fig S 5 a) for SO-spindle complexes, solitary SOs and solitary spindles.

**Supplementary Fig.6.**
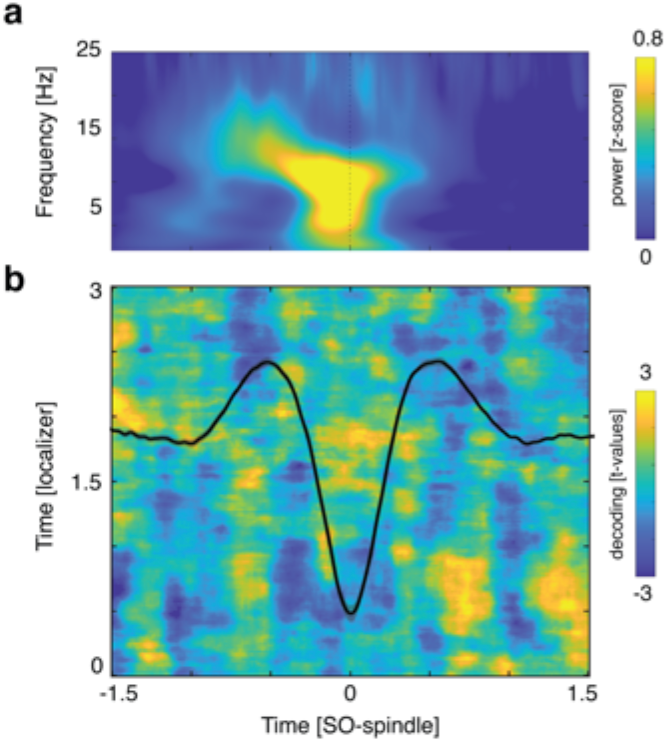

Slow spindle-SO locked memory reactivation

**(a)** Time frequency representation of all slow spindle - SO segments (z-scored across time; only positive values are displayed).

**(b)** When testing accuracy levels against chance at any localizer time x sleep time point, no significant cluster of above chance classification emerged (two-sided dependent-samples t-test; cluster with smallest p value: 0.67, cluster corrected across time). The black line illustrates the averaged EEG trace of all slow spindle - SO segments (electrode Fz).

**Supplementary Fig.7.**
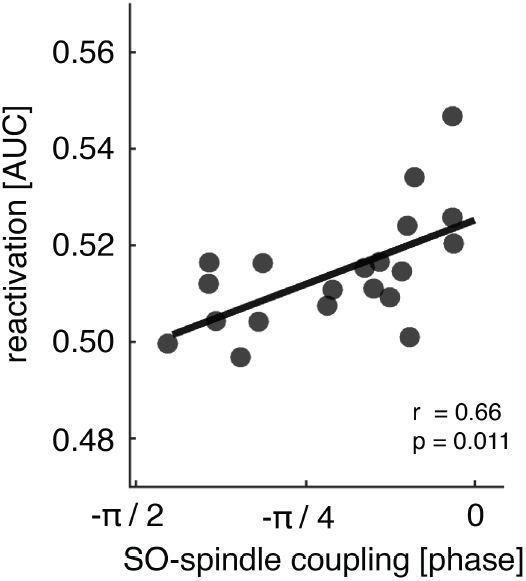
Precision of SO-spindle coupling correlates with reactivation strength. Circular-linear correlation analysis between the individual mean SO-spindle coupling phase (circles) and the mean reactivation strength (color coded) revealed a positive association (r = 0.66; p = 0.011).

**Supplementary Fig.8.**
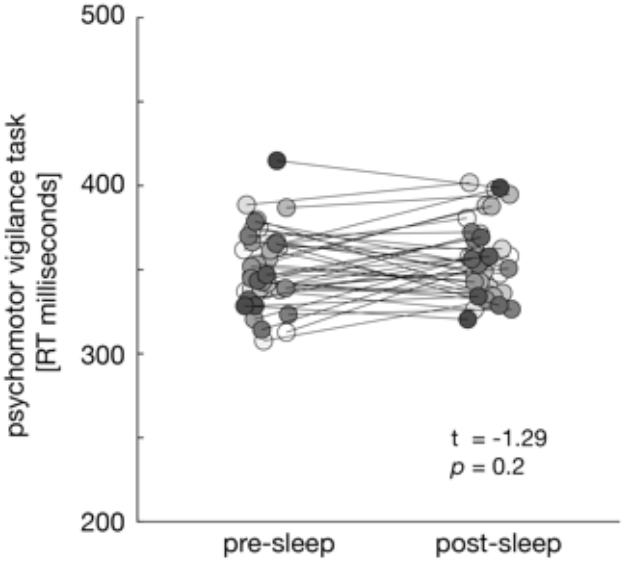
PVT results Before encoding and after the sleep period participants’ vigilance state was assessed using a modified version of the psychomotor vigilance task (“PVT”). During the pre-encoding task the mean reaction time was 350.6 ± 3.6 ms, while the average response time during the post-sleep PVT was 355.2 ± 3.6 ms. Reaction times did not differ between testing times (pre-encoding vs. post-sleep; t = −1.29, p = 0.20).

## Supplementary Tables

**Table 1:** Overview of memory performance

|  | Objects | Scenes | *t* | *P* |
| --- | --- | --- | --- | --- |
| Recognition [Hits] % |  |  |  |  |
| pre-sleep | 72.16 ± 4.16 | 70.91 ± 4.26 | 0.47 | 0.64 |
| post-sleep | 63.16 ± 4.19 | 63.58 ± 4.76 | -0.79 | 0.86 |
| post relative to pre | 87.41 ± 2.82 | 87.97 ± 2.88 | -0.16 | 0.85 |
| Recognition [Correct Rejections] % |  |  |  |  |
| pre-sleep | 90.00 ± 2.14 | 85.33 ± 4.48 | 1.13 | 0.27 |
| post-sleep | 88.00 ± 2.89 | 83.33 ± 5.22 | 0.98 | 0.33 |
| post relative to pre | 97.49 ± 1.75 | 97.33 ± 3.60 | 0.04 | 0.96 |
| Recognition [*d'*] |  |  |  |  |
| pre-sleep | 2.11 ± 0.14 | 2.02 ± 0.22 | 0.47 | 0.64 |
| post-sleep | 1.76 ± 0.19 | 1.69 ± 0.23 | 0.37 | 0.71 |
| Associative Memory % |  |  |  |  |
| pre-sleep | 49.16 ± 4.87 | 46.75 ± 3.08 | 1.13 | 0.41 |
| post-sleep | 40.08 ± 4.94 | 36.50 ± 4.46 | 1.11 | 0.28 |
| post relative to pre | 76.52 ± 5.27 | 72.61 ± 4.51 | 0.71 | 0.48 |
| Associative Memory [out of hits] % |  |  |  |  |
| pre-sleep | 64.90 ± 3.99 | 63.72 ± 5.20 | 0.33 | 0.73 |
| post-sleep | 59.39 ± 5.71 | 55.82 ± 5.47 | -1.03 | 0.31 |
| post relative to pre | 86.95 ± 5.52 | 82.55 ± 4.84 | 0.50 | 0.55 |

Associative memory % refers to the percentage of correctly recalled images (relative to the total number of stimuli), while associative memory [out of hits] refers to the percentage of recalled image exemplars out of correctly recognized verbs. Statistical differences between conditions (objects vs. scenes) were assessed using dependent samples *t*-tests (two-sided).

**Table 2:** Sleep characteristics. Data are means ± s.e.m. N1, N2: NREM sleep stages N1 & N2, SWS: slow-wave sleep, REM: rapid eye movement sleep, WASO: wake after sleep onset.

| Sleep stage [%] | Objects | Scenes | *t* | *P* |
| --- | --- | --- | --- | --- |
| N1 | 2.9 ± 1.6 | 13.5 ± 2.1 | -0.6 | 0.53 |
| N2 | 39.5 ± 2.6 | 48.1 ± 3.1 | -1.7 | 0.09 |
| SWS | 22.9 ± 3.3 | 19.4 ± 2.6 | -1.2 | 0.25 |
| REM | 21.9 ± 3.4 | 16.7 ± 2.8 | 1.4 | 0.15 |
| WASO | 2.3 ± 0.9 | 1.4 ± 0.7 | 0.7 | 0.49 |
| Total Sleep Time [min] | 102.6 ± 3.4 | 100.6 ± 2.9 | 0.6 | 0.53 |
| # spindles | 184.8 ± 12.9 | 188.9 ± 17.9 | -0.1 | 0.85 |
| Spindle density | 2.9 ± 0.1 | 2.82 ± 0.1 | 0.3 | 0.74 |
| Spindle duration | 0.81 ± 0.01 | 0.78 ± 0.01 | 1.3 | 0.19 |
| Spindle frequency | 14.02 ± 0.1 | 14.06 ± 0.1 | -0.2 | 0.81 |
| No SOs | 445.5 ± 31.2 | 474.1 ± 43.8 | -0.5 | 0.59 |
| SO density | 6.9 ± 0.25 | 7.1 ± 0.32 | -0.4 | 0.68 |
| SO duration | 1.38 ± 0.01 | 1.38 ± 0.01 | 0.5 | 0.61 |
| #SO – spindle comp.<br>range | 48.8 ± 3.8<br>[25-88] | 50.7 ± 4.9<br>[19-89] | -0.3 | 0.71 |

Statistical differences between conditions (objects vs. scenes) were assessed using dependent samples t-tests (two-sided).

**Table 3.**
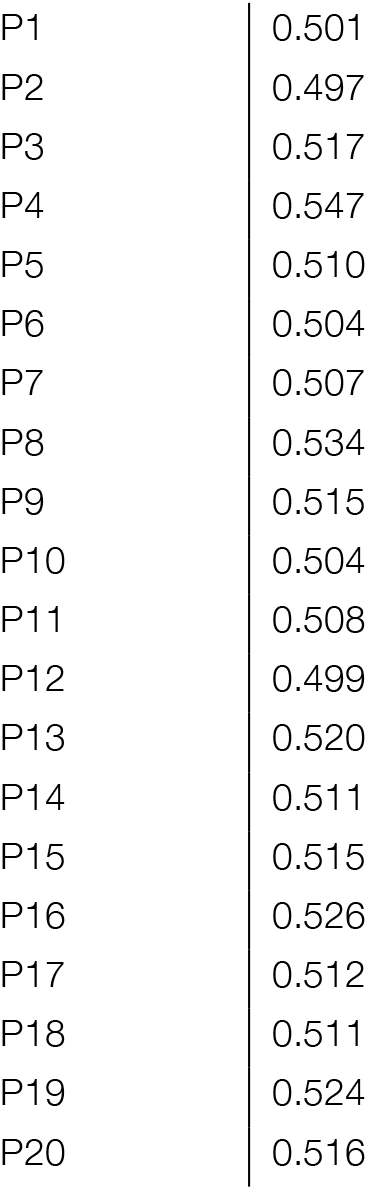
Mean decoding performance across the significant cluster of localizer – SO-spindle classification.

## Supplemental Notes

**Assessing the impact of trait-like characteristics on the interplay of memory reactivation with the preferred SO-spindle phase and the behavioral expressions of consolidation.**

Our results suggest that memory reactivation is linked to the preferred SO-spindle phase as well as to behavioral expressions of consolidation. However, SO-spindle coupling and consolidation might also be governed by other, trait-like participant characteristics. We thus examined - using hierarchical regressions - whether subjective sleep quality (as determined the Pittsburgh Sleep Quality Index (PSQI)) or circadian rhythm (derived from the Morningness-Eveningness Questionnaire (MEQ)) would account for memory performance or the preferred phase of SO-spindle coupling above and beyond reactivation strength during sleep.

In Step 1, reactivation strength explained 29.5% of the variance in memory performance (R^2^ = 0.295, F_1,18_= 7.51, P = 0.013). In Steps 2 and 3, neither subjective sleep quality (PSQI global score collapsed across the two sessions, range = 2-5 across participants), nor circadian rhythm (MEQ score collapsed across the two sessions, range = 31-66 across participants) explained significant amounts of additional variance (Step 2: ΔR^2^ = 0.001, ΔF_2,16_ = 0.014, P = 0.91; Step 3: ΔR^2^ < 0.01, ΔF_3,13_ < 0.01, P = 0.98). Similarly, using the preferred phase of SO-spindle coupling as the dependent variable, in Step 1 reactivation strength explained 37.2% of the variance (R^2^ = 0.372, F_1,18_= 10.65, P = 0.004). In Step 2 and 3, neither subjective sleep quality nor circadian rhythm explained significant amounts of additional variance (Step 2: ΔR^2^ = 0.07, ΔF_2,16_ = 0.18, P = 0.67; Step 3: ΔR^2^ = 0.01, ΔF_3,13_ = 0,30 P = 0.59). Please note that the distribution of the preferred phase values (all clustering between –π / 2 and 0 degrees) enabled us to add these circular data to the linear regression analysis. Nevertheless, to further test for potential associations between subjective sleep quality, circadian rhythm and the preferred phase of SO-spindle coupling, we administered additional circular-linear correlations. Neither sleep quality (rho = 0.04, p = 0.98) nor circadian rhythm (rho = 0.24, p = 0.47) correlated significantly with the preferred phase.

Together, although other trait-like characteristics not captured in the present analysis might have an influence, these results rule out that the link between memory reactivation and consolidation or the phase of SO-spindle coupling is driven by participants’ subjective sleep quality or circadian rhythm.

## References

1. Rasch, B. & Born, J. About Sleep’s Role in Memory. Physiol. Rev. 93, 681–766 (2013).

2. Walker, M. P. & Stickgold, R. Sleep-dependent learning and memory consolidation. Neuron 44, 121–33 (2004).

3. Diekelmann, S. & Born, J. The memory function of sleep. Nat. Rev. Neurosci. 11, 114–126 (2010).

4. Paller, K. A., Creery, J. D. & Schechtman, E. Memory and Sleep: How Sleep Cognition Can Change the Waking Mind for the Better. Annu. Rev. Psychol. 72, annurev-psych-010419-050815 (2021).

5. Buzsáki, G. The hippocampo-neocortical dialogue. Cerebral Cortex (1996). doi:10.1093/cercor/6.2.81

6. Marr, D. Simple memory: a theory for archicortex. Philos. Trans. R. Soc. Lond. B. Biol. Sci. 262, 23–81 (1971).

7. Sirota, A., Csicsvari, J., Buhl, D. & Buzsáki, G. Communication between neocortex and hippocampus during sleep in rodents. Proc. Natl. Acad. Sci. U. S. A. 100, 2065–2069 (2003).

8. Staresina, B. P. et al. Hierarchical nesting of slow oscillations, spindles and ripples in the human hippocampus during sleep. Nat. Neurosci. (2015). doi:10.1038/nn.4119

9. Maingret, N., Girardeau, G., Todorova, R., Goutierre, M. & Zugaro, M. Hippocampo-cortical coupling mediates memory consolidation during sleep. Nat. Neurosci. 19, 959–964 (2016).

10. Latchoumane, C.-F. V., Ngo, H.-V. V., Born, J. & Shin, H.-S. Thalamic Spindles Promote Memory Formation during Sleep through Triple Phase-Locking of Cortical, Thalamic, and Hippocampal Rhythms. Neuron 1–12 (2017). doi:10.1016/j.neuron.2017.06.025

11. Oyanedel, C. N., Durán, E., Niethard, N. & Marion Inostroza, |. R E S E A R C H R E P O R T Systems Neuroscience Temporal associations between sleep slow oscillations, spindles and ripples. Eur J Neurosci 00, 1–17 (2020).

12. Skelin, I., Kilianski, S. & McNaughton, B. L. Hippocampal coupling with cortical and subcortical structures in the context of memory consolidation. Neurobiol. Learn. Mem. 160, 21–31 (2019).

13. Steriade, M., McCormick, D. A. & Sejnowski, T. J. Thalamocortical oscillations in the sleeping and aroused brain. Science 262, 679–685 (1993).

14. Amzica, F. & Steriade, M. The functional significance of K-complexes. Sleep Med. Rev. 6, 139–149 (2002).

15. Timofeev, I. Neuronal plasticity and thalamocortical sleep and waking oscillations. in Progress in Brain Research 193, 121–144 (Elsevier B.V., 2011).

16. Isomura, Y. et al. Integration and Segregation of Activity in Entorhinal-Hippocampal Subregions by Neocortical Slow Oscillations. Neuron 52, 871–882 (2006).

17. Fernandez, L. M. J. & Lüthi, A. Sleep Spindles: Mechanisms and Functions. Physiological reviews 100, 805–868 (2020).

18. Mak-McCully, R. A. et al. Coordination of cortical and thalamic activity during non-REM sleep in humans. Nat. Commun. 8, 15499 (2017).

19. Rosanova, M. & Ulrich, D. Pattern-specific associative long-term potentiation induced by a sleep spindle-related spike train. J. Neurosci. 25, 9398–9405 (2005).

20. Seibt, J. et al. Cortical dendritic activity correlates with spindle-rich oscillations during sleep in rodents. Nat. Commun. 8, (2017).

21. Niethard, N., Ngo, H. V. V., Ehrlich, I. & Born, J. Cortical circuit activity underlying sleep slow oscillations and spindles. Proc. Natl. Acad. Sci. U. S. A. (2018). doi:10.1073/pnas.1805517115

22. Buzsáki, G. Hippocampal sharp wave-ripple: A cognitive biomarker for episodic memory and planning. Hippocampus (2015). doi:10.1002/hipo.22488

23. Joo, H. R. & Frank, L. M. The hippocampal sharp wave–ripple in memory retrieval for immediate use and consolidation. Nat. Rev. Neurosci. 19, 744–757 (2018).

24. Helfrich, R. F. et al. Bidirectional prefrontal-hippocampal dynamics organize information transfer during sleep in humans. Nat. Commun. (2019). doi:10.1038/s41467-019-11444-x

25. Ngo, H. V., Fell, J. & Staresina, B. Sleep spindles mediate hippocampal-neocortical coupling during long-duration ripples. Elife 9, (2020).

26. Clemens, Z. et al. Temporal coupling of parahippocampal ripples, sleep spindles and slow oscillations in humans. Brain 130, 2868–2878 (2007).

27. Jiang, X., Gonzalez-Martinez, J. & Halgren, E. Posterior Hippocampal Spindle Ripples Co-occur with Neocortical Theta Bursts and Downstates-Upstates, and Phase-Lock with Parietal Spindles during NREM Sleep in Humans. J. Neurosci. 39, 8949–8968 (2019).

28. Mikutta, C. et al. Phase‐amplitude coupling of sleep slow oscillatory and spindle activity correlates with overnight memory consolidation. J. Sleep Res. 28, (2019).

29. Zhang, J., Yetton, B., Whitehurst, L. N., Naji, M. & Mednick, S. C. The effect of zolpidem on memory consolidation over a night of sleep. Sleep (2020). doi:10.1093/sleep/zsaa084

30. Helfrich, R. F., Mander, B. A., Jagust, W. J., Knight, R. T. & Walker, M. P. Old Brains Come Uncoupled in Sleep: Slow Wave-Spindle Synchrony, Brain Atrophy, and Forgetting. Neuron 97, 221–230.e4 (2018).

31. Hahn, M. A., Heib, D., Schabus, M., Hoedlmoser, K. & Helfrich, R. F. Slow oscillation-spindle coupling predicts enhanced memory formation from childhood to adolescence. Elife 9, 1–21 (2020).

32. Muehlroth, B. E. et al. Precise Slow Oscillation–Spindle Coupling Promotes Memory Consolidation in Younger and Older Adults. Sci. Rep. 9, 1–15 (2019).

33. Kim, J., Gulati, T. & Ganguly, K. Competing Roles of Slow Oscillations and Delta Waves in Memory Consolidation versus Forgetting. Cell 179, 514–526.e13 (2019).

34. Oudiette, D. & Paller, K. A. Upgrading the sleeping brain with targeted memory reactivation. Trends Cogn. Sci. (2013).

35. Schreiner, T. & Staudigl, T. Electrophysiological signatures of memory reactivation in humans. Philosophical Transactions of the Royal Society B: Biological Sciences 375, (2020).

36. Cairney, S. A., Guttesen, A. á. V., El Marj, N. & Staresina, B. P. Memory Consolidation Is Linked to Spindle-Mediated Information Processing during Sleep. Current Biology (2018). doi:10.1016/j.cub.2018.01.087

37. Schreiner, T., Lehmann, M. & Rasch, B. Auditory feedback blocks memory benefits of cueing during sleep. Nat. Commun. 6, (2015).

38. Oyarzún, J. P., Morís, J., Luque, D., de Diego-Balaguer, R. & Fuentemilla, L. Targeted Memory Reactivation during Sleep Adaptively Promotes the Strengthening or Weakening of Overlapping Memories. J. Neurosci. 37, 7748–7758 (2017).

39. Epstein, R. & Kanwisher, N. A cortical representation of the local visual environment. Nature 392, 598–601 (1998).

40. Malach, R. et al. Object-related activity revealed by functional magnetic resonance imaging in human occipital cortex. Proc. Natl. Acad. Sci. U. S. A. 92, 8135–8139 (1995).

41. Antony, J. W., Ferreira, C. S., Norman, K. A. & Wimber, M. Retrieval as a Fast Route to Memory Consolidation. Trends Cogn. Sci. xx, 1–4 (2017).

42. Roediger, H. L. & Karpicke, J. D. Test-enhanced learning: Taking memory tests improves long-term retention. Psychol. Sci. 17, 249–255 (2006).

43. Macmillan, N. A. & Creelman, C. D. Detection Theory: A User’s Guide. Detection Theory A users guide Standort:, (Cambridge University Press, 2005).

44. Ngo, H. V. V, Martinetz, T., Born, J. & Mölle, M. Auditory closed-loop stimulation of the sleep slow oscillation enhances memory. Neuron 78, 545–553 (2013).

45. Davachi, L. Item, context and relational episodic encoding in humans. Current Opinion in Neurobiology 16, 693–700 (2006).

46. Klinzing, J. G., Niethard, N. & Born, J. Mechanisms of systems memory consolidation during sleep. Nature Neuroscience (2019). doi:10.1038/s41593-019-0467-3

47. Schönauer, M. et al. Decoding material-specific memory reprocessing during sleep in humans. Nat. Commun. 8, 15404 (2017).

48. Bergmann, T. O., Mölle, M., Diedrichs, J., Born, J. & Siebner, H. R. Sleep spindle-related reactivation of category-specific cortical regions after learning face-scene associations. Neuroimage 59, 2733–42 (2012).

49. Rasch, B., Büchel, C., Gais, S. & Born, J. Odor cues during slow-wave sleep prompt declarative memory consolidation. Science (80-.). 315, 1426–1429 (2007).

50. Rudoy, J. D., Voss, J. L., Westerberg, C. E. & Paller, K. A. Strengthening individual memories by reactivating them during sleep. Science (80-.). 326, 1079 (2009).

51. Schreiner, T., Doeller, C. F., Jensen, O., Rasch, B. & Staudigl, T. Theta Phase-Coordinated Memory Reactivation Reoccurs in a Slow-Oscillatory Rhythm during NREM Sleep. Cell Rep. 25, 296–301 (2018).

52. Göldi, M., van Poppel, E. A. M., Rasch, B. & Schreiner, T. Increased neuronal signatures of targeted memory reactivation during slow-wave up states. Sci. Rep. 9, 2715 (2019).

53. Wang, B. et al. Targeted Memory Reactivation during Sleep Elicits Neural Signals Related to Learning Content. J. Neurosci. 39, 6728–6736 (2019).

54. Bar, E. et al. Local Targeted Memory Reactivation in Human Sleep. Curr. Biol. 30, 1435–1446.e5 (2020).

55. Schechtman, E. et al. Multiple memories can be simultaneously reactivated during sleep as effectively as a single memory. bioRxiv 662015 (2019). doi:10.1101/662015

56. Swanson, R. A., Levenstein, D., McClain, K., Tingley, D. & Buzsáki, G. Variable specificity of memory trace reactivation during hippocampal sharp wave ripples. Current Opinion in Behavioral Sciences (2020). doi:10.1016/j.cobeha.2020.02.008

57. Belal, S. et al. Identification of memory reactivation during sleep by EEG classification. Neuroimage 176, 203–214 (2018).

58. Schapiro, A. C. et al. Sleep Benefits Memory for Semantic Category Structure while Preserving Exemplar-Specific Information. Sci. Rep. 7, (2017).

59. Drosopoulos, S., Schulze, C., Fischer, S. & Born, J. Sleep’s function in the spontaneous recovery and consolidation of memories. J. Exp. Psychol. Gen. 136, 169–183 (2007).

60. Cairney, S. A., Lindsey, S., Sobczak, J. M., Paller, K. A. & Gaskell, G. The Benefits of Targeted Memory Reactivation for Consolidation in Sleep Are Contingent on Memory Accuracy and Direct Cue-Memory Associations.: Sleep and Memory Reactivation. Sleep 44, (2016).

61. Creery, J. D., Oudiette, D., Antony, J. W. & Paller, K. A. Targeted Memory Reactivation during Sleep Depends on Prior Learning. Sleep 38, 755–63 (2015).

62. Petzka, M., Charest, I., Balanos, G. & Staresina, B. Does sleep-dependent consolidation favour weak memories? (2020). doi:10.31234/osf.io/q4wnv

63. Rothschild, G. The transformation of multi-sensory experiences into memories during sleep. Neurobiol. Learn. Mem. (2019). doi:10.1016/j.nlm.2018.03.019

64. Rothschild, G., Eban, E. & Frank, L. M. A cortical-hippocampal-cortical loop of information processing during memory consolidation. Nat. Neurosci. (2017). doi:10.1038/nn.4457

65. Navarrete, M., Valderrama, M. & Lewis, P. A. The role of slow-wave sleep rhythms in the cortical-hippocampal loop for memory consolidation. Current Opinion in Behavioral Sciences 32, 102–110 (2020).

66. Sejnowski, T. J. & Destexhe, A. Why do we sleep? Brain Res. 886, 208–223 (2000).

67. Antony, J. W. et al. Sleep Spindle Refractoriness Segregates Periods of Memory Reactivation. Curr. Biol. 28, 1736–1743.e4 (2018).

68. Ngo, H.-V. V. et al. Driving Sleep Slow Oscillations by Auditory Closed-Loop Stimulation--A Self-Limiting Process. J. Neurosci. 35, 6630–6638 (2015).

69. Buysse, D. J., Reynolds, C. F., Monk, T. H., Berman, S. R. & Kupfer, D. J. The Pittsburgh sleep quality index: A new instrument for psychiatric practice and research. Psychiatry Res. (1989). doi:10.1016/0165-1781(89)90047-4

70. Horne, J. A. & Ostberg, O. A self assessment questionnaire to determine Morningness Eveningness in human circadian rhythms. Int. J. Chronobiol. (1976).

71. Dinges, D. F. & Powell, J. W. Microcomputer analyses of performance on a portable, simple visual RT task during sustained operations. Behav. Res. Methods, Instruments, Comput. (1985). doi:10.3758/BF03200977

72. Konkle, T., Brady, T. F., Alvarez, G. A. & Oliva, A. Conceptual distinctiveness supports detailed visual long-term memory for real-world objects. J. Exp. Psychol. Gen. 139, 558–578 (2010).

73. Brainard, D. H. The Psychophysics Toolbox. Spat. Vis. 10, 433–436 (1997).

74. Iber, C., Ancoli-Israel, S., Chesson, A. & Quan, S. The AASM Manual for the Scoring of Sleep and Associated Events: Rules, Terminology, and Technical Specification. American Academy of Sleep Medicine (2007).

75. Oostenveld, R., Fries, P., Maris, E. & Schoffelen, J. M. FieldTrip: Open source software for advanced analysis of MEG, EEG, and invasive electrophysiological data. Comput. Intell. Neurosci. 2011, (2011).

76. Demanuele, C. et al. Coordination of slow waves with sleep spindles predicts sleep-dependent memory consolidation in schizophrenia. Sleep (2017). doi:10.1093/sleep/zsw013

77. Jung, T.-P. et al. Extended ICA removes artifacts from electroencephalographic recordings. in Advances in Neural Information Processing Systems 10, 894–900 (1998).

78. Silber, M. H. et al. The visual scoring of sleep in adults. Journal of Clinical Sleep Medicine (2007). doi:10.5664/jcsm.26814

79. Cox, R. & Fell, J. Analyzing human sleep EEG: A methodological primer with code implementation. (2020). doi:10.1016/j.smrv.2020.101353

80. De Gennaro, L. & Ferrara, M. Sleep spindles: an overview. Sleep Med. Rev. 7, 423–440 (2003).

81. Berens, P. CircStat : A MATLAB Toolbox for Circular Statistics. J. Stat. Softw. 31, 1–21 (2009).

82. Treder, M. S. MVPA-Light: A Classification and Regression Toolbox for Multi-Dimensional Data. Front. Neurosci. 14, (2020).

83. Jackson, J. E. A Use’s Guide to Principal Components. (John Wiley & Sons, Inc., 1991). doi:10.1002/0471725331

84. Jiang, Y. & Guo, P. Regularization versus dimension reduction, which is better? in Lecture Notes in Computer Science (including subseries Lecture Notes in Artificial Intelligence and Lecture Notes in Bioinformatics) 4492 LNCS, 474–482 (Springer Verlag, 2007).

85. Grootswagers, T., Wardle, S. G. & Carlson, T. A. Decoding dynamic brain patterns from evoked responses: A tutorial on multivariate pattern analysis applied to time series neuroimaging data. J. Cogn. Neurosci. 29, 677–697 (2017).

86. Pinheiro-Chagas, P., Piazza, M. & Dehaene, S. Decoding the processing stages of mental arithmetic with magnetoencephalography. (2019). doi:10.1016/j.cortex.2018.07.018

87. Sankaran, N., Thompson, W. F., Carlile, S. & Carlson, T. A. Decoding the dynamic representation of musical pitch from human brain activity. Sci. Rep. 8, 1–9 (2018).

88. Lemm, S., Blankertz, B., Dickhaus, T. & Müller, K. R. Introduction to machine learning for brain imaging. Neuroimage 56, 387–399 (2011).

89. King, J. R. & Dehaene, S. Characterizing the dynamics of mental representations: The temporal generalization method. Trends in Cognitive Sciences 18, 203–210 (2014).

90. Maris, E. & Oostenveld, R. Nonparametric statistical testing of EEG- and MEG-data. J. Neurosci. Methods 164, 177–190 (2007).

